# MORC proteins regulate transcription factor binding by mediating chromatin compaction in active chromatin regions

**DOI:** 10.1101/2022.11.01.514783

**Authors:** Zhenhui Zhong, Yan Xue, C. Jake Harris, Ming Wang, Zheng Li, Yunqing Ke, Yasaman Jami-Alahmadi, Suhua Feng, James A. Wohlschlegel, Steven E. Jacobsen

## Abstract

**Background:** The Microrchidia (MORC) proteins are a family of evolutionarily conserved GHKL-type ATPases involved in chromatin compaction and gene silencing. Arabidopsis MORC proteins act in the RNA-directed DNA methylation (RdDM) pathway, where they act as molecular tethers to ensure the efficient establishment of RdDM and *de novo* gene silencing. However, MORC proteins also have RdDM-independent functions; although, their underlying mechanisms are unknown.

**Results:** In this study, we examined regions of MORC binding where RdDM does not occur in order to shed light on the RdDM-independent functions of MORC proteins. We found that MORC proteins compact chromatin and reduce DNA accessibility to transcription factors (TFs), thereby repressing gene expression. We also found that MORC-mediated repression of gene expression was particularly important under conditions of stress. We showed that MORC proteins regulate TFs through either direct or indirect interactions, and these TFs can in some cases regulate their own transcription, resulting in feedforward loops.

**Conclusions:** Our findings provide insights into the molecular mechanisms of MORC-mediated chromatin compaction and transcription regulation.

## Background

The MORC proteins are a family of highly conserved GHKL-type ATPases involved in gene silencing and chromatin compaction [1]. In *C. elegans*, MORC-1 can compact DNA through a through topological entrapment [2], while in humans, MORC2 is recruited by the human silencing hub (HUSH) complex for H3K9me3 deposition, chromatin compaction, and gene silencing [3]. In mice, MORC1 is involved in germline transposon silencing [4], and MORC3 is essential for transposon silencing in embryonic stem cells [5].

The Arabidopsis genome encodes six MORC proteins: MORC1, 2, 4, 5, 6, and 7 (MORC3 being a pseudogene) [6]. These six proteins are functionally redundant, but colocalize with sites of RNA-directed DNA methylation (RdDM) genome-wide [7], where they are critical for establishing efficient RdDM and *de novo* gene silencing [7]. MORC7, when tethered to DNA using an artificial zinc finger, can target RdDM to ectopic sites. MORC7 is also required for the silencing of a newly integrated *FWA* transgene [7]. MORC proteins also act downstream of DNA methylation to suppress gene expression, and are also involved in plant immunity — protecting against potential pathogens by interacting with plant resistance (R) proteins [8,9]. However, the molecular mechanisms underlying these RdDM-independent functions remain unknown. We previously observed MORC binding sites where RdDM does not occur (MORC-unique sites) [7]; and by studying these sites, we aim to shed light on the mechanisms underlying the RdDM-independent functions of MORC proteins.

TOPLESS (TPL) and LEUNIG (LUG) are both Grocho (Gro)/TLE-type transcriptional co-repressors in plants. They are characterized by a conserved glutamine-rich C-terminal domain and an N-terminal WD-repeat domain [10]. The glutamine-rich domain participates in protein oligomerization, and the WD-repeat domain interacts with downstream transcriptional regulators [10]. The functional counterpart of the Gro/TLE family of proteins in yeast, Tup1, was originally identified as a co-repressor that occupied the binding sites of transcriptional activators [11,12].

However, evidence now shows that Tup1 can switch from a co-repressor to a co-activator in response to stress, and is required for the activation of certain genes related to the stress response [12,13].

Here, we use MORC-unique sites to study the RdDM-independent functions of MORC proteins. We show that MORC proteins compact chromatin and reduce DNA accessibility to TFs, thereby repressing the transcription of stress-responsive genes.

## Results

### MORC proteins bind to active chromatin regions devoid of RdDM

We previously reported that approximately 80% of MORC7 binding regions overlap with sites of RdDM [7]. MORC7 is recruited to these sites by the RdDM machinery, where it then facilitates the efficiency of the RdDM pathway. However, the remaining 20% of MORC7 binding sites are devoid of RdDM, as evidenced by a lack of Pol V occupancy [7]. The mechanisms underlying the function of MORC7 within these RdDM-depleted regions remain unknown.

Mouse MORC3 recognizes and localizes to regions of H3K4me3-marked chromatin through its CW domain [14]; however, Arabidopsis MORCs do not contain CW domains. To determine whether Arabidopsis MORCs co-localize with specific chromatin features, we used the ChromHMM method to investigate correlations between MORC7 and several well-characterized chromatin features (H3K9ac, H3K27ac, H4K16ac, H3K4me1, H3K4me3, H3K36me2, H3K36me3, H3K9me2, H3K27me3, Pol II, and Pol V). We analyzed chromatin states using a similar method as previously reported [15] but also included Pol V ChIP-seq data. We found 13 different chromatin states (Supplementary Fig. 1). MORC7 showed a strong correlation with Pol V (a known indicator of RdDM sites), which was consistent with our previous findings (State 11, Supplementary Fig. 1). Chromatin state 12 included sites enriched with MORC7 but depleted of Pol V — indicative of MORC7-unique regions. We did not observe enrichment of histone marks in these MORC7-unique regions (Supplementary Fig. 1).

Within these MORC7-unique regions, we identified two subgroups: MORC7A and MORC7B. The ChIP-seq data for MORC7, Pol V, ATAC-seq and transposable element (TE) density indicated that MORC7A was within a region of high chromatin accessibility and low TE density (Fig. 1a, b). Consistent with the ChromHMM analysis, MORC7A displayed low levels of histone occupancy and histone modification, although its flanking regions were enriched for active histone modifications. This suggests that MORC7A is located within an active chromatin compartment between genes (Fig. 1b).

**Fig 1.**
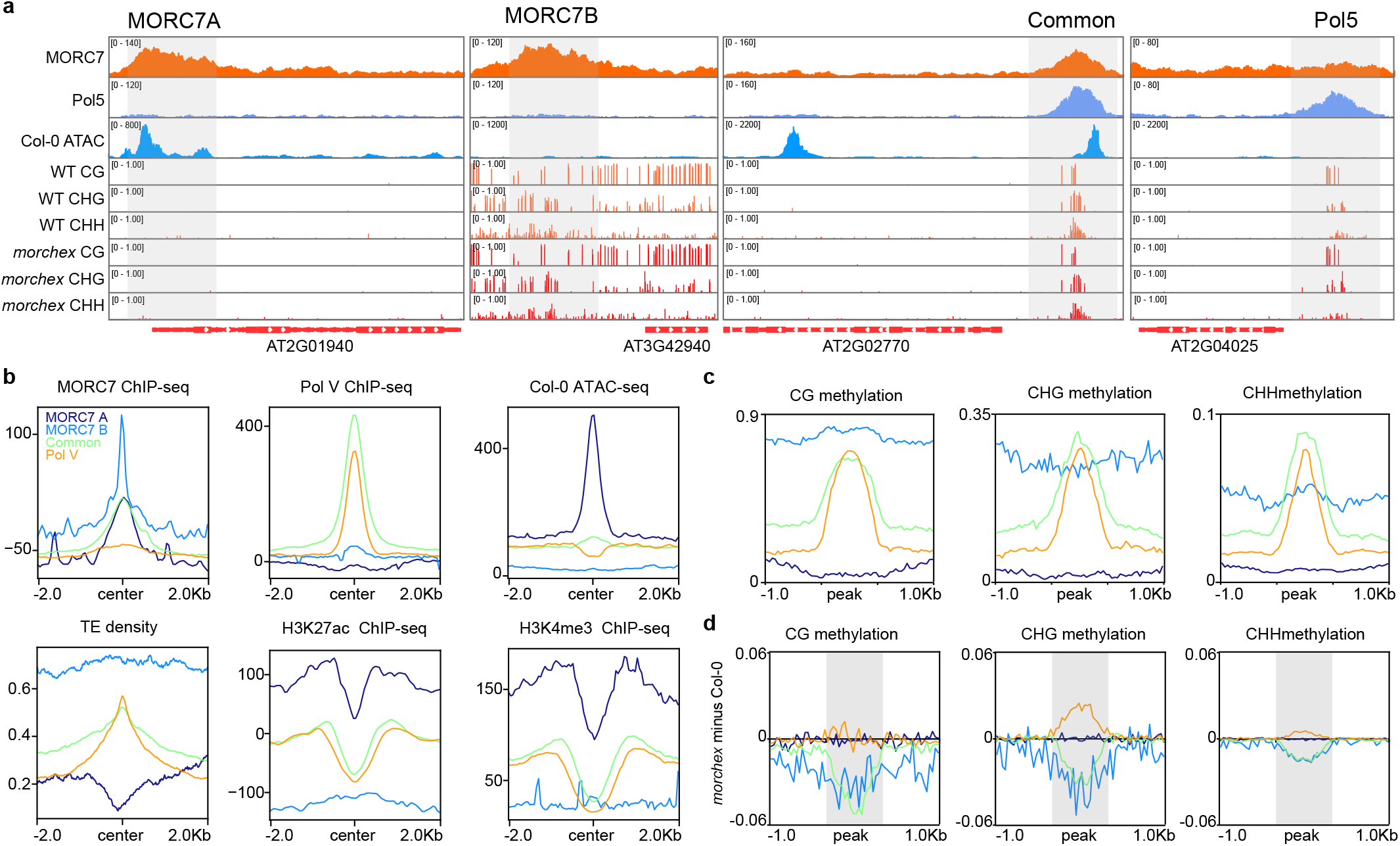
MORC7 binds to regions devoid of RdDM. **a**. Screenshots of ChIP-seq data for MORC7A-unique, MORC7B-unique, MORC7-Pol V Common, and Pol V-unique regions. **b**. Metaplots of ChIP-seq data for MORC7, Pol V, H3K27ac, H3K4me3, ATAC-seq, and transposable element (TE) density over regions of MORC7A-unique, MORC7B-unique, MORC7-Pol V Common, and Pol V-unique. **c**. Metaplot showing methylation levels of CG, CHG, and CHH, over regions of MORC7A-unique, MORC7B-B unique, MORC7-Pol V Common, and Pol V-unique. **D**. Metaplot showing methylation levels of CG, CHG, and CHH methylation changes (*morchex* minus WT) over regions of MORC7A-unique, MORC7B-unique, MORC7-Pol V Common, and Pol V-unique.

MORC7B contained a high density of TE with no apparent active histone marks, reflective of its heterochromatic location (Fig. 1b). We found that MORC7A regions had low levels of DNA methylation, while MORC7B regions had high levels of methylation (Fig. 1c, d). These results suggest that MORC7 binds to active and deep heterochromatic regions of DNA, where RdDM does not occur, suggesting that it regulates gene expression at these sites through RdDM-independent mechanisms.

### MORC7 preferentially binds to the promoters of TFs

The genomic distribution enrichment data showed enrichment of MORC7A peaks over promoters (Fig. 2a), but no enrichment of MORC7B peaks — consistent with their deep heterochromatic localization. The functional annotation of the genes proximal to MORC7A suggested that they were enriched with TFs (Table 1). The Arabidopsis genome encodes approximately 1491 TF genes (5.5% of the genome) [16]. Of the genes proximal to MORC7A, 23% were TFs (p-value = 3.12E-36); these included PHYTOCHROME INTERACTING FACTOR (PIF), ethylene and auxin-responsive transcriptional factors and Myb transcriptional factors (Supplementary Table 1). This enrichment was more significant than enrichment for MORC7-Pol V common (p-value = 0.001) and Pol V-unique (p-value = 1.6E-5).

**Fig 2.**
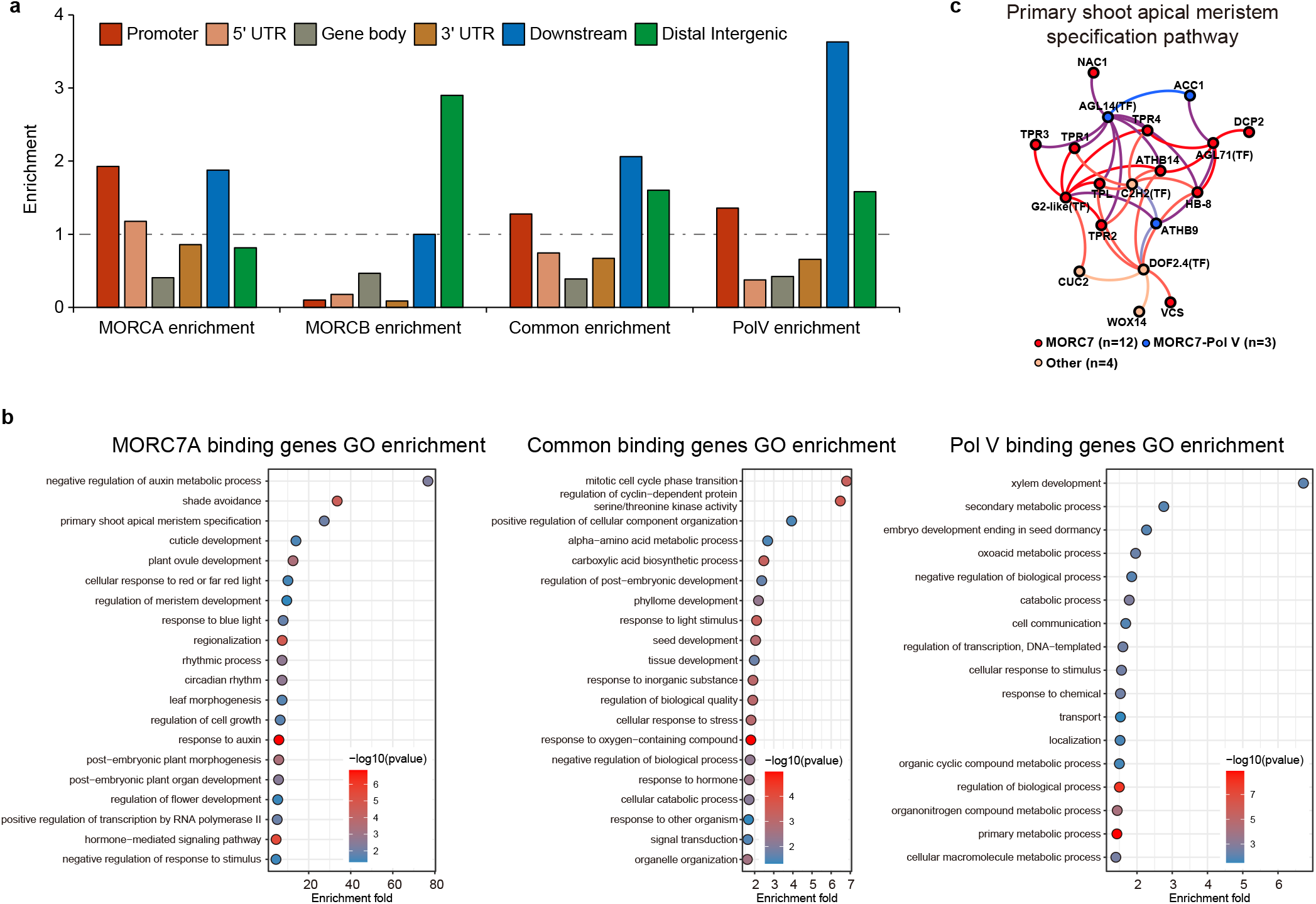
MORC7-unique regions preferentially localize to the promoter regions of TFs. **a**. Genomic distribution enrichment data for MORC7A-unique, MORC7B-unique, MORC7-Pol V Common, and Pol V-unique regions. **b**. Gene ontology enrichment data for the proximal genes of MORC7A, MORC7-Pol V Common, and Pol V-unique regions. **c**. MORC7 and Pol V binding on promoters of genes in the primary shoot apical meristem specification pathway.

**Table 1.** Number of TFs among the proximal genes of MORC7A-unique, MORC7B-unique, MORC7-Pol V Common, and Pol V-unique regions.

| Peak | TF | non TF | Total | Percentage | p-value |
| --- | --- | --- | --- | --- | --- |
| MORC7A | 99 | 326 | 425 | 23.29% | 3.12E-36 |
| MORC7B | 6 | 27 | 33 | 18.18% | 0.002 |
| MORC7-Common | 114 | 1475 | 1589 | 7.17% | 0.001 |
| Pol V | 98 | 1080 | 1178 | 8.32% | 1.60E-05 |
| Whole genome | 1491 | 25681 | 27172 | 5.49% | NA |

Gene Ontology (GO) term analysis of genes proximal to MORC7A showed an enrichment of negative regulation of auxin metabolic process (~80 fold), shade avoidance (~30 fold), and the primary shoot apical meristem specification pathway (~30 fold) (Fig. 2b). The primary shoot apical meristem specification pathway (GO0010072) is responsible for the growth of all post-embryonic, above-ground plant structures [17]. In Arabidopsis, this pathway includes several topless-related genes [17]. Interestingly, we found that MORC7 specifically bound to 12 of the 19 genes in this pathway (Fig. 2c, Supplementary Fig. 2), and co-localized with Pol V at an additional three. We show examples of MORC7 enrichment over the promoter regions for the four TOPLESS genes in Supplementary Fig. 2.

### MORC7 closely co-localizes with some TFs

To investigate the protein interaction network of MORC7 with chromatin, we re-analyzed previously published crosslinked IP-MS data of MORC7 [7]. We identified 494 proteins (FDR < 0.05, FC > 2) that interacted with MORC7 (Fig. 3a), and found that many of these were involved in either chromatin-related pathways or development (Fig. 3b). We also identified 68 TFs from the MORC7 interacting proteins (68/494, p = 7.89E-12) (Supplementary Table 2). To further test whether MORC7 co-localizes with TFs, we obtained binding site information for 200 TFs from the DNA Affinity Purification and sequencing (DAP-seq) database [18], and performed pairwise peak overlap analysis with MORC7 peaks. We found that MORC7A showed stronger co-localization with TFs compared to MORC7B, MORC7-Pol V common and Pol V-unique regions (Fig. 3c). This indicates that MORC7A peaks are associated with TF binding sites. We also re-analyzed three factors in particular because published ChIP-seq data was available, PIF4 [19], ARF6 [20], and TPR1 [21]. Metaplot analysis showed that with ChIP-seq data for MORC7A-unique, MORC7B-unique, MORC7-Pol V Common, and Pol V-unique regions showed the strongest co-localization with MORC7A (Fig. 3d, Supplementary Fig. 3).

**Fig. 3.**
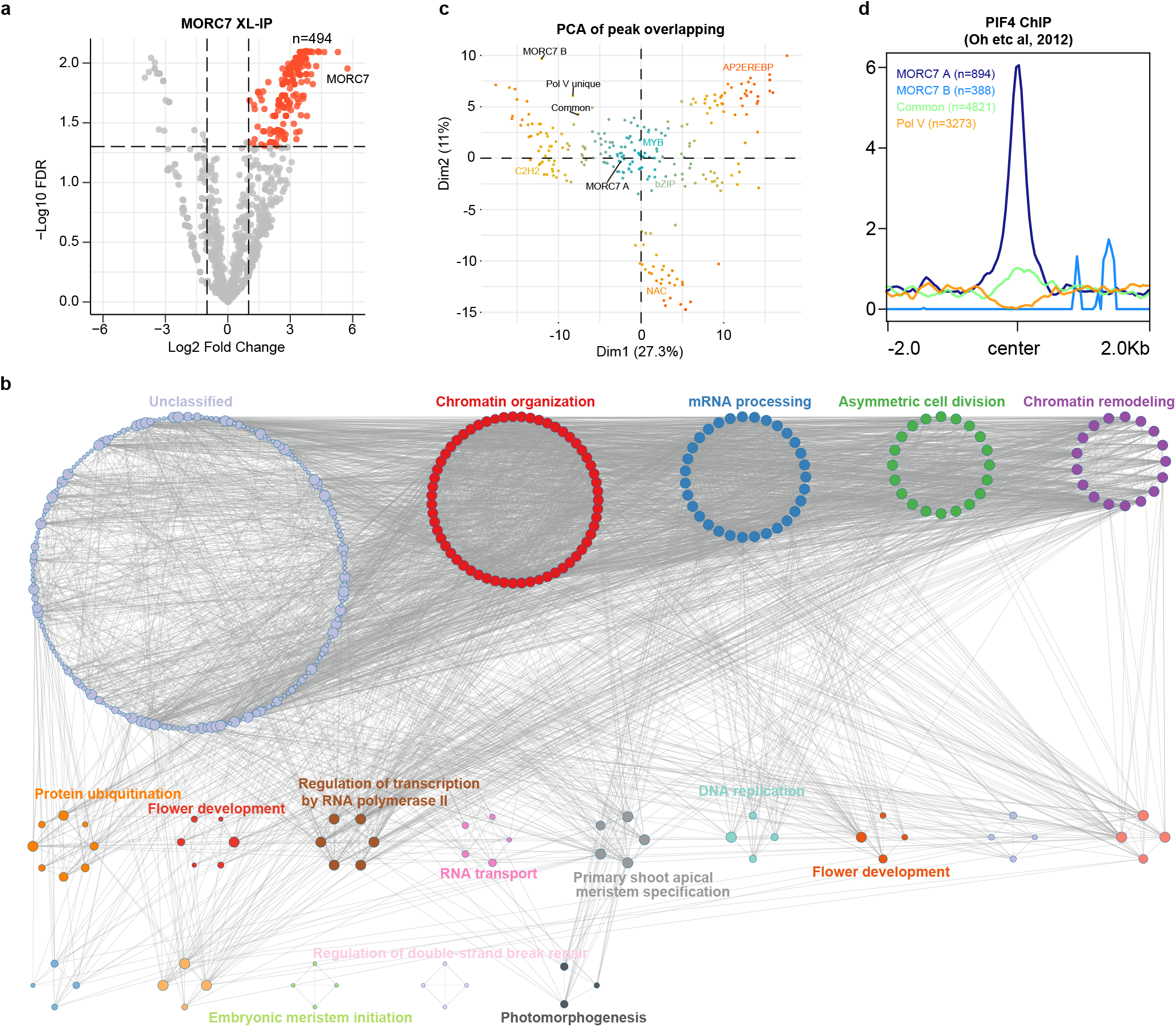
MORC7 associates with some TFs. **a**. Volcano plot showing proteins that have significant interactions with MORC7, as detected by crosslinked IP-MS. **b**. Protein-protein interaction networks of MORC7. **c**. A graph showing the degree of overlap between the DAP-seq peaks of approximately 200 TFs with MORC7A-unique, MORC1B-unique, MORC7-Pol V Common, and Pol V unique regions. **d**. Metaplot of PIF4 ChIP-seq data [19] over MORC7A-unique, MORC7B-unique, MORC7-Pol V Common, and Pol V unique regions.

### MORC7 influences TF binding through chromatin compaction

To understand how MORC7 affects chromatin conformation, we performed an Assay for Transposase-Accessible Chromatin with high-throughput sequencing (ATAC-seq) in *morc4 morc7*, *morc6*, and *morc hextuple* (*morchex*, in which all functional MORCs are knocked out) mutants [6]. We plotted ATAC-seq data across the four groups, and found that MORC7A, MORC7B and MORC7-Pol V common showed greater chromatin accessibility changes in the mutants, particularly over the MORC7A regions (Fig. 4a, b). This phenotype is consistently observed in *morc4 morc7, morc6* and *morche*x — with *morchex* showing the most pronounced phenotype (Fig. 4a). Interestingly, for Pol V-unique sites, DNA compaction was not reduced, but actually became slightly increased (Fig. 4a). Consistently, we also observed an increase in DNA methylation for Pol V-unique sites in the mutants (Fig. 1d). This suggests that Pol V may be redistributed from the MORC7-Pol V common sites to Pol V-unique sites in the absence of MORC proteins. This is consistent with our previous findings that suggested MORC proteins function as molecular tethers to facilitate the recruitment of RdDM components [7].

**Fig. 4.**
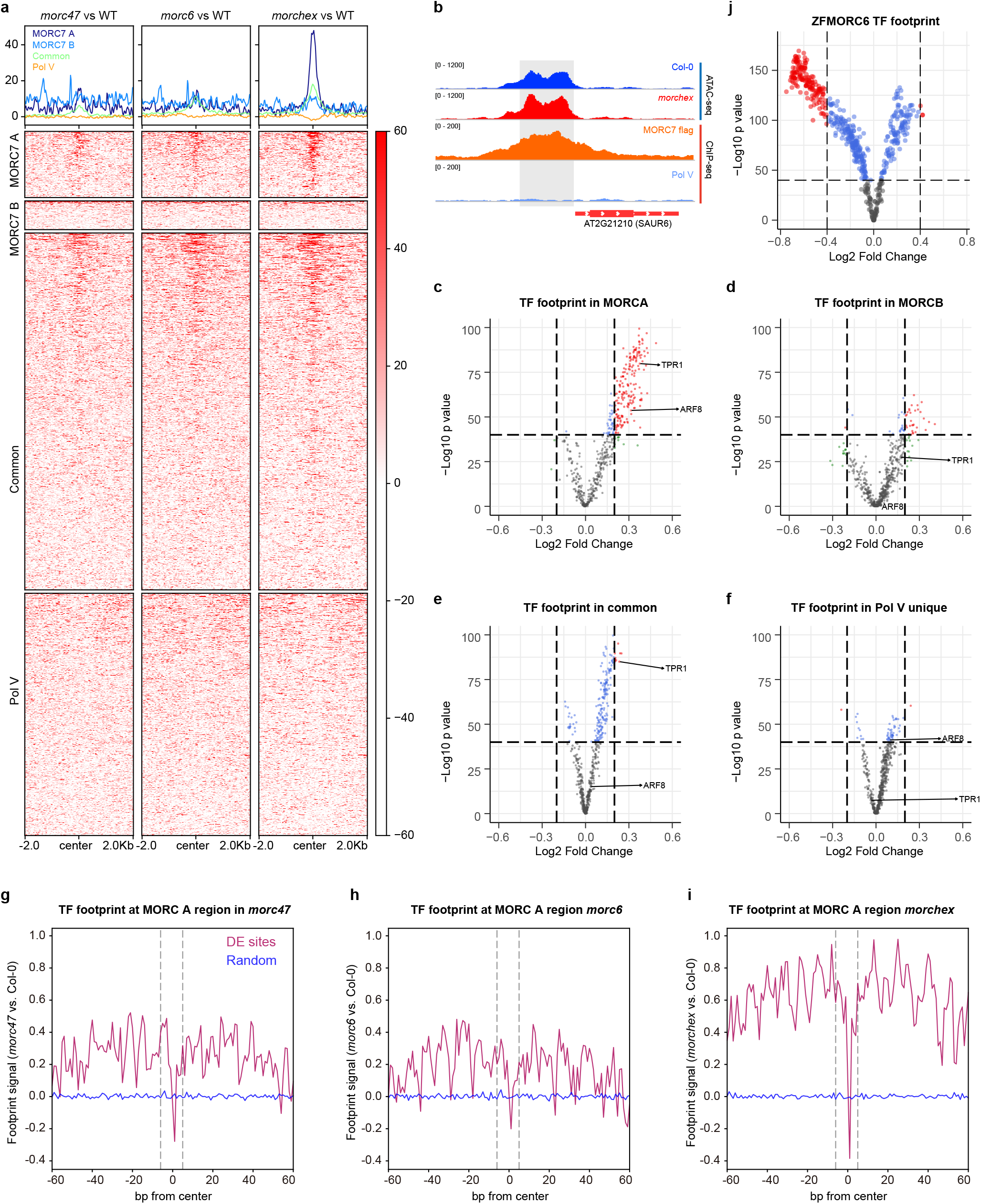
MORC proteins influence TF binding through chromatin compaction. **a**. Metaplot showing chromatin accessibility changes in MORC7A-unique, MORC7B-unique, MORC7-Pol V Common, and Pol V-unique regions profiled by ATAC-seq. **b**. A representative screenshot showing higher chromatin accessibility at the promoter of SAUR6 in the *morchex* mutant. **c**. Volcano plot showing changes in TF footprints in MORC7A regions, comparing *morchex* and wild type. **d**. Volcano plot showing changes in TF footprints in MORC7B regions, comparing *morchex* and wild type. **e**. Volcano plot showing changes in TF footprints in MORC7-Pol V Common regions, comparing *morchex* and wild type. **f**. Volcano plot showing changes in TF footprints at Pol V-unique regions, comparing *morchex* and wild type. **g**. Metaplot showing TF footprint changes for MORC7A-unique regions in the *morc4morc7* mutant. **h**. Metaplot showing TF footprint changes for MORC7A-unique regions in the *morc6* mutant. **i**. Metaplot showing TF footprint changes for MORC7A-unique regions in the *morchex* mutant. **j**. Volcano plot showing TF changes for ZF off-target sites, comparing ZF-MORC6 and *fwa-4* plants.

To examine whether MORC-mediated DNA compaction affects TFs, we analyzed the ATAC-seq data for TF footprints. When a TF binds to DNA, it inhibits the integration of DNA by Tn5 transposes, causing the binding motif to exhibit lower DNA accessibility, and the flanking regions to exhibit higher DNA accessibility [22]. The footprints of 572 TFs downloaded from JASPAR were analyzed in the *morc4 morc7*, *morc6* and *morchex* mutants [23]. Many TFs showed substantially stronger apparent binding within the MORC7A regions in the mutants. There were some increases in binding within the MORC7B regions (although to a lesser degree than in MORC7A regions) (Fig. 4c, d), while TF binding over RdDM sites was largely unaffected (Fig. 4e, f). The metaplot of ATAC-seq signals over the TF binding sites for the MORC7A regions confirmed that these TFs have stronger apparent binding in *morc4 morc7, morc6* and *morche*x mutants – with *morchex* showing the strongest binding changes, and the random control regions showing no differences (Fig. 4g-i).

We previously showed that targeting either MORC7 or MORC6 ectopically in the *fwa-4* epiallele background using ZF108 can trigger the silencing of *FWA* [7,24]. In addition to the *FWA* locus, ZF108 can also bind thousands of off-target sites [24]. These off-target sites are preferentially localized to promoter regions, and therefore provide an excellent opportunity to test whether the presence of MORC proteins can affect TF binding. We compared TF footprints between ZF-MORC6 and *fwa-4* and found a substantial decrease for many of the TF footprints in ZF-MORC6 plants. This supports the hypothesis that MORC proteins affect TF binding (Fig. 4j). Together, these results suggest that MORCs inhibit TF binding by altering chromatin accessibility.

### MORC influences gene expression downstream of the TFs

To understand whether MORC proteins regulate gene expression, we performed RNA-seq with the *morchex* mutant. As MORC7A co-localizes strongly with PIF4 (Fig. 3d) – a central regulator in temperature signaling [25] – we applied heat treatment to the *morchex* mutant. We first compared the expression of genes proximal to MORC7A peaks in wild type (WT) and *morchex* mutant without treatment. This showed that the genes proximal to MORC7A were slightly up-regulated in the *morchex* mutant without treatment (Fig. 5a), including the TFs SEP3, PIF4, ARF6/8, TPR1, LUG and SEU (Fig. 5b). After heat treatment, *morchex* displayed a stronger response compared to the WT, with significantly more upregulated genes (Fig. 5c, d). Genes proximal to MORC7A were enriched in shoot apical meristem specification pathways, and consistently, we observed stronger upregulation of these genes in *morchex* after heat treatment (Supplementary Fig. 4).

**Fig. 5.**
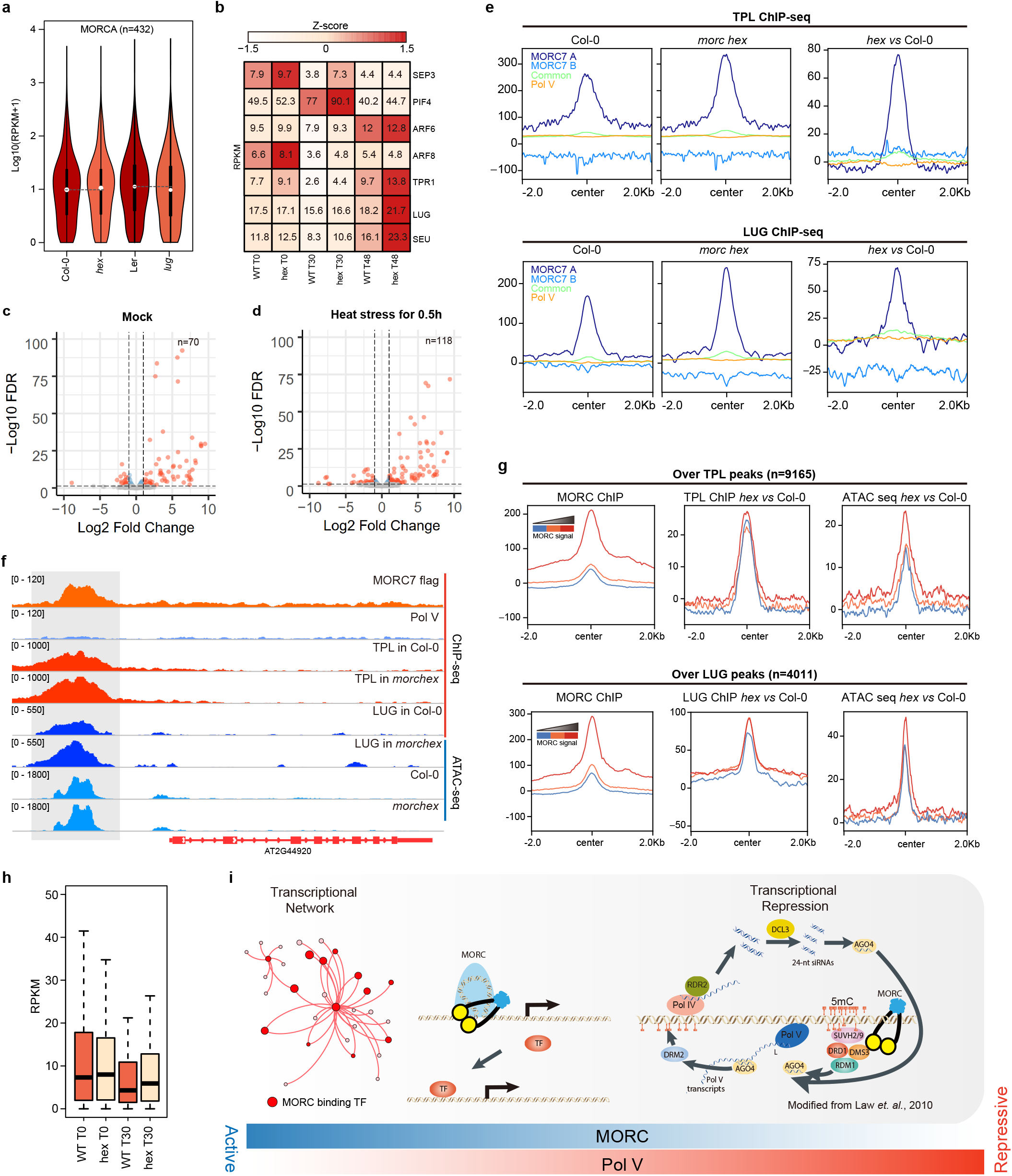
MORC influence TF binding through chromatin compaction. Violin plot showing expression levels of genes proximal to MORC7A with Col-0, *morchex* mutant, Ler (wild type background for *lug* mutant), and *lug* mutant. **b**. Expression levels of transcriptional factors: SEP3, PIF4, ARF6/8, TPR1, LUG and SEU (TFs with MORC7A peaks in their promoter regions), with Col-0 and *morchex* mutants following heat treatment. **c**. Transcriptomic changes of *morchex* mutants under normal conditions. **d**. Transcriptomic changes of *morchex* mutants after 30 minutes of heat treatment. **e**. TPL and LUG binding over MORC7A-unique, MORC7B-unique, MORC7-Pol V Common, and Pol V unique regions. **f**. A representative screenshot showing increased binding of TPL and LUG on MORC7A-unique regions in the *morchex* mutant. **g**. Correlation of TPL/LUG binding and ATAC-seq alterations with MORC7 binding intensity in *morchex* mutant. **h**. Boxplot showing the expression levels of genes directly regulated by LUG in Col-0 and *morchex* mutants following heat treatment for 30 minutes (T30). **i**. A proposed model of the RdDM-independent functions of MORC proteins.

To confirm that MORC proteins affect TF binding, and to understand how they affect downstream gene expression, we selected TFs TOPLESS (TPL) and LEUNIG (LUG) for ChIP-seq analysis, because they were present in the MORC7 IP-MS data [7]. We expressed TPL and LUG fused with a 3XFLAG-tag in both WT and *morchex*. Consistent with the TF footprint analysis, both TPL and LUG displayed stronger binding at MORC7A regions, while only a slight increase in binding was noted for the MORC7-Pol V co-binding sites in *morchex* (Fig. 5e, f) *—* consistent with an increase in chromatin accessibility in *morchex* (Fig. 5g; Supplementary Fig. 5). We ranked TPL and LUG binding sites based on the MORC ChIP-seq signals and divided them into three groups; high, middle and low. Overall, in the *morchex* mutant, we observed increased TPL and LUG binding, as well as increased chromatin accessibility across the regions with stronger MORC7 signals (Fig. 5g). We found that MORC7A-bound genes were downregulated in the *lug* mutant, suggesting that LUG may facilitate expression of these genes (Fig. 5a). Using ChIP-seq data together with RNA-seq data in the *lug* mutant, we identified 95 genes that appeared to be directly regulated by LUG (Supplementary Table 3). We found that these LUG-regulated genes were upregulated in *morchex*, particularly after heat treatment (Fig. 5h).

## Discussion

We previously reported that MORC proteins are localized to sites of RdDM throughout the genome, and function as molecular tethers to facilitate the efficient establishment of RdDM [7]. We showed that this RdDM-related function of MORC proteins is critical for *de novo* transgene silencing [7,24]; however, this model does not explain other functions of MORC proteins. For example, MORC1 and MORC6 were shown to work downstream of DNA methylation to repress the expression of both the endogenous *SDC* gene and an *SDC* transgene, as well as other DNA methylated targets in the genome [9]. In addition, *morc* mutants display various disease phenotypes; for example, Kang et al. [8] reported that *morc1* is susceptible to Turnip Crinkle Virus (TRV), while Harris et al. [6] reported that *morchex* is susceptible to the *Hyaloperonospora arabidopsidis* (Hpa) strain, Emwa1. However, the molecular mechanisms underlying the additional functions of the MORC proteins remain unknown.

Here, we investigated the function of MORC7 in regions where RdDM does not occur, particularly those near genes where no DNA methylation is present. We found that MORC proteins reduce chromatin accessibility within these regions. Previous *in vitro* studies showed that *C. elegans* MORC1 homodimers can topologically entrap and condense DNA through further oligomerization of MORC1 proteins [2]. In addition, Arabidopsis *morc* mutants display pericentromeric heterochromatin decondensation [9], which takes place with minimal losses of DNA methylation throughout the genome. This indicates that MORC proteins contribute to chromatin compaction independently of DNA methylation [9]. We show here that MORC proteins reduce chromatin accessibility in methylation-free promoter regions of DNA, which may explain their mechanism for methylation-independent gene regulation. We suggest that Arabidopsis MORCs may use a similar mechanism of chromatin compaction to that of *C. elegans* MORC1 – compacting chromatin by topological entrapment, thereby reducing its accessibility to TFs.

Plant MORCs have been implicated in plant pathogen responses. MORCs promote resistance in some plant species and inhibit defense responses in others [8]. Upregulation of protein-coding genes was previously shown in *morc4 morc7*; although, the underlying mechanisms of this was unknown [6]. Here, we report that MORC proteins regulate gene expression by compacting chromatin in promoter regions, thereby preventing access by TFs. In addition, MORCs preferentially bound to the promoter regions of TFs, contributing to their regulation, and our crosslinked IP-MS data suggested that MORCs are located in close proximity to some TFs. Interestingly, we observed that many of these TFs bind to their own promoters – suggesting egulation by a feedforward loop, which may amplify the effects of MORCs on transcriptional networks.

Finally, we showed that MORC proteins are important for the regulation of gene expression, particularly under stress conditions. We also found altered expression of heat-responsive genes in *morchex*. Like with its role in plant pathogen defense response, it seems likely that the role of MORCs in stress responses relates to its chromatin compaction of promoter regions and affects in TF networks.

## Conclusions

MORC proteins have a broad binding spectrum in the genome, and appear to participate in at least three separate processes. They co-localize to sites of RdDM, facilitating efficient DNA methylation establishment [7], they are needed to repress DNA methylated areas of pericentromeric heterochromatin in a DNA methylation independent manner [9], and they co-localize with TFs in unmethylated promoter regions, regulating TF binding and gene expression by altering chromatin accessibility (Fig. 5i). Although it seems likely that MORC act in each of these processes by topologically entrapping DNA, there are likely mechanistic differences that can explain the localization and function of MORCs in these three different epigenetic environments in the genome.

## Methods

### Plant materials and growth conditions

All plants in this study were grown in standard greenhouse conditions (22 — 25 °C, 16 hrs light/8 hrs dark). The following plant materials were used in this study: *morchex* consisting of *morc1-2* (SAIL_893_B06), *morc2-1* (SALK_072774C), *morc4-1* (SALK_051729), *morc5-1* (SALK_049050C), *morc6-3* (GABI_599B06), and *morc7-1* (SALK_051729). For heat treatments, plants were grown under 37 °C for 0.5 hours and put back to normal temperature for 48 hours for recovery.

### Epitope-tagged transgenic lines

Full-length genomic DNA fragments, including native promoter sequences, were cloned into pENTR/D vectors (Invitrogen), followed by modified destination vectors carrying 3xFLAG with LR Clonase (Invitrogen). All primers used in this study are available in Supplementary Table 4.

### Nuclei extraction and ATAC-seq library preparation

The nuclei collection process from inflorescence and meristem tissues was performed in accordance with previously described methods [26,27]. Freshly isolated nuclei were used for ATAC-seq, as described elsewhere [28]. Inflorescence tissues were collected for extraction of nuclei as follows: 5g (approximately) of inflorescence tissue was collected and immediately transferred into the ice-cold grinding buffer (300mM sucrose, 20mM Tris pH 8, 5mM MgCl2, 5mM KCl, 0.2% Triton X-100, 5mM β-mercaptoethanol, and 35% glycerol); the samples were then ground with Omni International General Laboratory Homogenizer at 4°C, and filtered through a two-layer Miracloth using a 40-μm nylon mesh Cell Strainer (Fisher). Samples were spin filtered for 10 min at 3,000 *g*, the supernatant was discarded, and the pellet was resuspended with 25ml of grinding buffer using a Dounce homogenizer. The wash step was performed twice in total. Nuclei were then resuspended in 0.5ml of freezing buffer (50mM Tris pH 8, 5mM MgCl2, 20% glycerol, and 5mM β-mercaptoethanol). Nuclei were then subjected to a transposition reaction with Tn5 (Illumina). For the transposition reaction, 25μl of 2 x DMF (66mM Tris-acetate pH 7.8, 132mM K-Acetate, 20mM Mg-Acetate, and 32% DMF) was mixed with 2.5μl Tn5 and 22.5μl nuclei suspension at 37°C for 30 min. The transposed DNA fragments were then purified with ChIP DNA Clean & Concentrator Kit (Zymo). Libraries were prepared with Phusion High-Fidelity DNA Polymerase (NEB), in a system containing: 12.5μl 2 x Phusion, 1.25μl 10mM Ad1 primer, 1.25μl 10mM Ad2 primer, 4μl ddH2O, and 6μl purified transposed DNA fragments. The ATAC-seq libraries were sequenced on the NovaSeq 6000 platform (Illumina).

### RNA-seq library preparation

Total RNAs were extracted from 100mg (approximately) of flower buds using TRIzol and the Direct-zol RNA Miniprep kit (Zymo, R2050). Sequencing libraries were prepared using the TruSeq Stranded mRNA Library Prep kit (Illumina), according to the manufacturer’s instructions, and sequenced on a NovaSeq 6000 sequencer (Illumina).

### ChIP-seq library preparation

10g of inflorescence and meristem tissues were used for ChIP-seq. ChIP assays were performed as has been described previously [29]. Briefly, 2-4g of flower tissue was collected from 4 — 5-week-old plants, and ground with liquid nitrogen. 1% formaldehyde containing a nuclei isolation buffer was used to fix the chromatin for ten minutes. Freshly prepared glycine was then used to terminate the crossing reaction. Shearing was performed via Bioruptor Plus (Diagenode), and immunoprecipitations with antibodies were performed overnight at 4°C. Magnetic Protein A and Protein G Dynabeads (Invitrogen) were added and incubated at 4°C for two hours. The reverse crosslink was performed overnight at 65°C. The protein-DNA mix was then treated with Protease K (Invitrogen) at 45°C for four hours. The DNA was purified and precipitated with 3M Sodium Acetate (Invitrogen), glycoBlue (Invitrogen) and Ethanol overnight at −20°C. The precipitated DNA was then used for library preparation using the Ovation Ultra Low System V2 kit (NuGEN), which was then sequenced using an Illumina NovaSeq sequencer. The anti-FLAG M2 (Sigma) antibody was used in this study. Libraries were prepared using the NuGen Ovation Ultra Low System V2 kit, in accordance with the manufacturer’s instructions.

### ATAC-seq analysis

ATAC-seq read adaptors were removed using trim_galore. The reads were then mapped to the Arabidopsis thaliana reference genome, TAIR10, using Bowtie2 (−X 2000 −m 1) [30]. Reads of chloroplast and mitochondrial DNA were filtered out and duplicate reads were removed using Samtools [31]. ATAC-Seq open chromatin peaks of each replicate were called using MACS2 with parameters of −p 0.01 --nomodel --shift −100 --extsize 200. Consensus sets of chromatin peaks for all samples were merged by bedtools (v2.26.0) intersect allowing a distance of 10 base pairs [32]. Following this, edgeR was used to define significant changes between peaks [Fold Change, (FC) > 2 and False Discovery Rate, (FDR) < 0.05] [33]. ATAC-seq peak distributions were annotated using ChIPseeker [34]. TF footprints were analyzed by TOBIAS [22] with 572 plant TF motifs downloaded from JASPAR (http://jaspar.genereg.net/) [23].

### RNA-seq analysis

Cleaned short reads were aligned to the reference genome, TAIR10, by Bowtie2 (v2.1.0) [30]. Expression abundance was then calculated by RSEM using the default parameters [35]. Heatmaps were visualized using the R package pheatmap. Differential expression analysis was conducted using edgeR [33]. A threshold of p-value < 0.05 and Fold Change > 2 were used to decide whether there were any significant differences in expression between samples.

### ChIP-seq analysis

ChIP-seq data was aligned to the TAIR10 reference genome with Bowtie2 (v2.1.0) [30], only including uniquely mapped reads without any mismatches. Duplicated reads were removed by Samtools. ChIP-seq peaks were called by MACS2 (v2.1.1) and annotated using ChIPseeker [34]. Differential peaks were called by the bdgdiff function in MACS2 [36]. ChIP-seq data metaplots were plotted by deeptools (v2.5.1) [37]. Correlation of MORC7 with ChIP-seq data was conducted with ChromHMM [38]. H3K9ac, H3K27ac, H4K16ac, H3K4me1, H3K4me3, H3K36me2, H3K36me3, H3K9me2, H3K27me3, Pol II, and Pol V, as published previously, were included in this analysis [26,39–44] (Supplementary Table 5).

### Whole-genome bisulfite sequencing (BS-seq) analysis

Previously published whole-genome bisulfite sequencing data for *morc*-mutants and wild type was reanalyzed [6]. Briefly, Trim_galore (http://www.bioinformatics.babraham.ac.uk/projects/trim_galore/) was used to trim adapters. BS-seq reads were aligned to the TAIR10 reference genome by BSMAP (v2.90), allowing two mismatches and one best hit (-v 2 -w 1) [45]. Reads with three or more consecutive CHH sites were considered to be unconverted reads and were filtered out. DNA methylation levels were defined as #C/ (#C + #T).

## Supplementary Information

The online version contains supplementary material available at xxxx.

## Acknowledgements

We thank members of Jacobsen laboratory for the helpful discussion. We are grateful to M. Akhavan and the UCLA BSCRC High Throughput BioSequencing Core for their technical assistance. We also thank Life Science Editors (https://www.lifescienceeditors.com/) for editing assistance.

## Peer review information

xxx was the primary editor of this article and managed its editorial process and peer review in collaboration with the rest of the editorial team.

## Review history

The review history is available as Additional file x.

## Availability of data and materials

Data supporting the findings of this work are available within the paper and its Supplementary Information files. All high-throughput sequencing data generated in this study are accessible at NCBI’s Gene Expression Omnibus (GEO) via GEO Series accession number GSE212801 (https://www.ncbi.nlm.nih.gov/geo/query/acc.cgi?acc=GSE212801). The customized codes used in this study are available upon reasonable request.

## Authors’ contributions

ZZ, YX, and SEJ conceived the study. ZZ, YX, and CJH performed experiments assisted by MW, ZL, and YK. SF performed high throughput sequencing. YJA and JAW performed IP-MS. ZZ, YX, and SEJ wrote the manuscript with help from all authors. All authors approved the final version of the manuscript and agree on the content and conclusions.

## Declarations

### Ethics approval and consent to participate

Not applicable.

### Consent for publication

Not applicable.

## Supplementary figures

**Supplementary Fig. 1 Chromatin states of MORC7**. ChIP-seq analysis was performed for H4K16ac, H3K4me3, H3K27ac, H3K9ac, H3Ac, H3K36me3, H3K4me1, H3K36me2, H3K9me2, H3K27me3, Pol II, Pol V, and MORC7.

**Supplementary Fig. 2** Examples showing MORC7 enrichment over the promoter regions of the TOPLESS genes.

**Supplementary Fig. 3. MORC7 associates with some TFs. a**. Metaplot of ARF6 ChIP-seq data [20] over MORC7A-unique, MORC7B-unique, MORC7-Pol V Common, and Pol V-unique regions. Metaplot of TPR1 ChIP-seq data [21] over MORC7A-unique, MORC7B-unique, MORC7-Pol V Common, and Pol V-unique regions. **c**. A screenshot showing MORC7A co-localization with TPR1.

**Supplementary Fig. 4** Expression levels of genes in the primary shoot apical meristem specification pathway, with and without heat treatment, in Col-0 and *morchex* mutants.

## Supplementary Tables

**Supplementary Table 1** Genes proximal to MORC7A-unique, MORC7B-unique, MORC7-Pol V Common, and Pol V-unique peaks.

**Supplementary Table 2** List of MORC7 interacting proteins.

**Supplementary Table 3** Expression level of LUG directly regulated genes in Ler and lug mutant.

**Supplementary Table 4** Primers used in this study.

**Supplementary Table 5** Published ChIP-seq data used for ChromHMM states analysis.

