## Supplementary figures for "MORC proteins regulate transcription factor binding by mediating chromatin compaction in active chromatin regions"

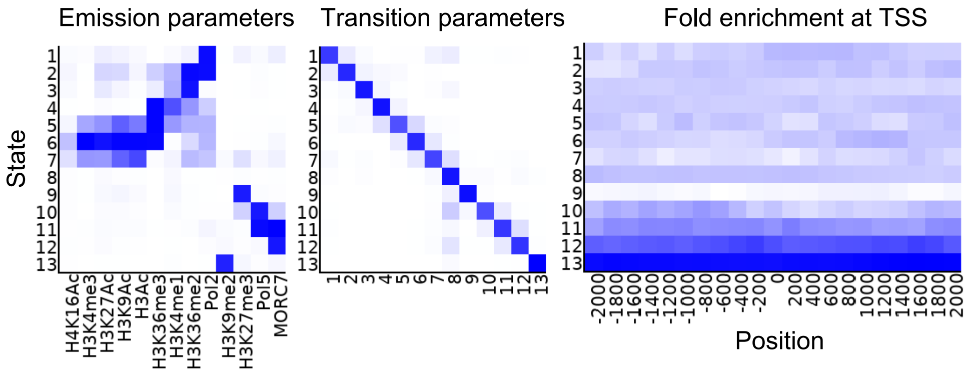


**Supplementary Fig. 1 Chromatin states of MORC7.** ChIP-seq analysis was performed for H4K16ac, H3K4me3, H3K27ac, H3K9ac, H3Ac, H3K36me3, H3K4me1, H3K36me2, H3K9me2, H3K27me3, Pol II, Pol V, and MORC7.

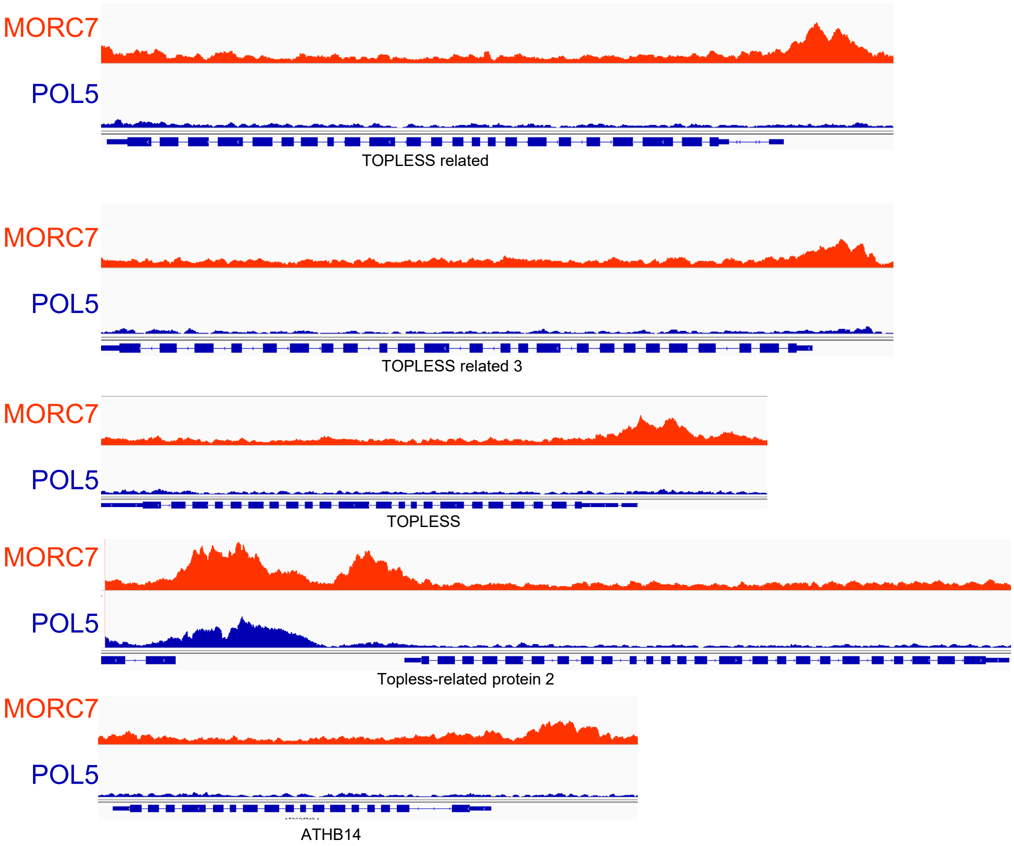


**Supplementary Fig. 2** Examples showing MORC7 enrichment over the promoter regions of the TOPLESS genes.

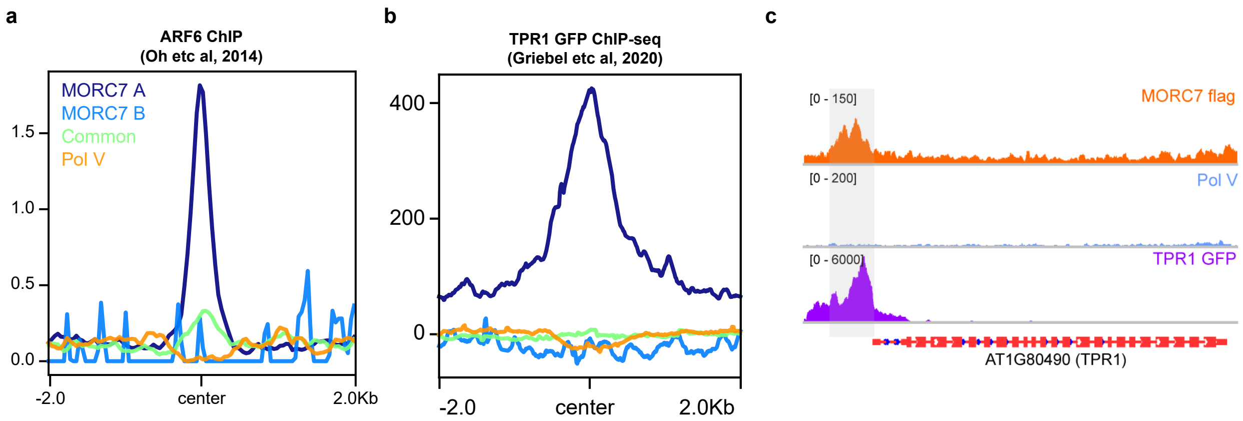


**Supplementary Fig. 3. MORC7 associates with some TFs. a.** Metaplot of ARF6 ChIP-seq data [20] over MORC7A-unique, MORC7B-unique, MORC7-Pol V Common, and Pol V-unique regions. **b.** Metaplot of TPR1 ChIP-seq data [21] over MORC7A-unique, MORC7B-unique, MORC7-Pol V Common, and Pol V-unique regions. **c.** A screenshot showing MORC7A co-localization with TPR1.

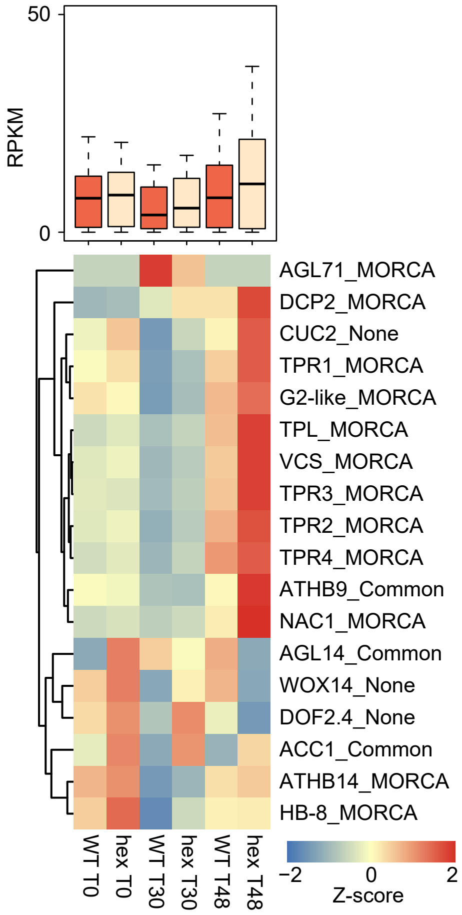


**Supplement Fig. 4** Expression levels of genes in the primary shoot apical meristem specification pathway, with and without heat treatment, in Col-0 and *morchex* mutants.
