## Supplementary table 5 for "MORC proteins regulate transcription factor binding by mediating chromatin compaction in active chromatin regions"

**Supplementary Table 5** Published ChIP-seq data used for ChromHMM states analysis.

| **Data** | **SRA link** |
| --- | --- |
| H3K27me3 | SRR10905142 [1] |
| H3K27ac | SRR1509479 [2] |
| H3K9me2 | SRR6410845 [3] |
| H3K4me3 | SRR10905140 [1] |
| H3K9ac | SRR4733909 [4] |
| H4K16ac | SRR6364625 [5] |
| H3K4me1 | SRR8742329 [6] |
| H3K36me2 | SRX13085623 |
| H3K36me3 | SRX13085627 |
| Pol II | SRX13085631 |
| Pol V | SRR5681073 [7] |
